# TRILL: Orchestrating Modular Deep-Learning Workflows for Democratized, Scalable Protein Analysis and Engineering

**DOI:** 10.1101/2023.10.24.563881

**Authors:** Zachary A Martinez, Richard M. Murray, Matt W. Thomson

**Affiliations:** Division of Biology and Bioengineering, California Institute of Technology, Pasadena, CA 91125

## Abstract

Deep-learning models have been rapidly adopted by many fields, partly due to the deluge of data humanity has amassed. In particular, the petabases of biological sequencing data enable the unsupervised training of protein language models that learn the “language of life.” However, due to their prohibitive size and complexity, contemporary deep-learning models are often unwieldy, especially for scientists with limited machine learning backgrounds. TRILL (**TR**aining and **I**nference using the **L**anguage of **L**ife) is a platform for creative protein design and discovery. Leveraging several state-of-the-art models such as ESM-2, DiffDock, and RFDiffusion, TRILL allows researchers to generate novel proteins, predict 3-D structures, extract high-dimensional representations of proteins, functionally classify proteins and more. What sets TRILL apart is its ability to enable complex pipelines by chaining together models and effectively merging the capabilities of different models to achieve a sum greater than its individual parts. Whether using Google Colab with one GPU or a supercomputer with hundreds, TRILL allows scientists to effectively utilize models with millions to billions of parameters by using optimized training strategies such as ZeRO-Offload and distributed data parallel. Therefore, TRILL not only bridges the gap between complex deep-learning models and their practical application in the field of biology, but also simplifies the orchestration of these models into comprehensive workflows, democratizing access to powerful methods. Documentation: **https://trill.readthedocs.io/en/latest/home.html**.

## Introduction

Even though the first genome was sequenced in 1977 [1], there are now more than 36 petabytes of sequencing data on the NCBI Sequence Read Archive [2]. This scientific feat has enabled scientists to utilize the breadth of nature for their research; whether it be for revealing unculturable organisms that live in extreme environments [3], to discovering novel antimicrobial peptides [4] or finding new ways to engineer genomes [5]. While there are seemingly endless amounts of sequencing data, actually leveraging this treasure-trove of biological novelty is a non-trivial task. Classical methods for comparing biological sequences usually involve pairwise comparisons [6] or by using hidden-markov models (HMMs) [7]. While these methods rely on evolutionary relationships to link related sequences through homology, machine learning based methods have shown success for functional comparisons without needing shared ancestry. For example, researchers have been able to predict whether a given protein is a cell-penetrating peptide, regardless of actual homology [8]. These predictions were enabled by extracting amino acid frequencies and biochemical properties for each protein and using this data to train random-forest classifiers. However, these methods tend to rely heavily on feature selection, which not only requires background knowledge but also introduce potential bias by focusing on select features.

Deep learning methods, such as recurrent neural networks, have proven to be able to extract functional information of proteins from sequence alone [9]. However, the application of natural language processing (NLP) methods to protein sequences has truly led to breakthroughs for state-of-the-art in-silico protein analysis. At the heart of these breakthroughs is the Transformer, a deep-learning architecture first published in 2017, which uses self-attention to effectively capture long-range dependencies and intricate patterns in protein sequences that were previously challenging for traditional methods to discern [10]. Analogous to using words and sentences to train typical large language models (LLMs), Transformer-based models such as ESM-2 use amino acids and protein sequences [11] to learn the “grammar” of life. These protein language models (PLMs) learn in a self-supervised manner, where the model attempts to predict the identity of a random 15% of the amino acids per sequence using the unmasked portions. ESM-2 was pre-trained on the masked language training task with 65 million unique protein sequences from UniRef. After this extensive training, scientists are able to use these pre-trained models to extract high-dimensional representations for their proteins of interest. These vectors can then be used for clustering, regression, classification and other downstream tasks for functional comparisons [12]. However useful deep-learning based approaches are, large PLMs are unwieldy, with ESM-2 parameter sizes ranging from millions to billions of parameters, much too big for many GPUs. The sheer size of these types of models limit widespread adoption in the biological community, where most end-users are not able to efficiently wield these powerful tools due to hardware constraints. While tools such as bio_embeddings [13] and ColabFold/Design [14] exist, no current software allows for both embedding and fine-tuning with these large PLMs at scale. Furthermore, end-to-end analyses for proteins often require chaining together methods, including other types of models like denoising diffusion probabilistic models and graph based message passing neural networks.

Recent advancements in the field have provided us with novel methods and examples for protein engineering. Despite the excitement, we identified a significant gap in the availability of computational platforms designed for agile, multifaceted experimentation in protein analysis. Motivated by this unaddressed need and eager to use these methods ourselves, we initiated the development of TRILL (**TR**aining and **I**nference using the **L**anguage of **L**ife). TRILL provides users with an expansive toolkit that encompasses a wide array of functionalities, from de-novo protein design and 3D structure prediction to simulating protein-ligand interactions and assessing protein properties. The platform’s user-friendly and flexible interface not only facilitates efficient model utilization for researchers of varying backgrounds but also reduces technical barriers. This enables a more focused and declarative approach to advancing protein science. A distinguishing feature of TRILL is its capability to compose complex workflows, allowing for more comprehensive analyses. By integrating these features, TRILL serves as a “toolbox,” aiming to become a go-to resource for both biologists and bioengineers engaged in protein-related research.

### The TRILL Framework

TRILL serves as a comprehensive platform designed to democratize access to state-of-the-art deep-learning models and methods. The framework eliminates the need for specialized computational skills, thereby lowering the entry barrier for end-users. At present, TRILL offers a rich repository of deep-learning models specifically trained for protein design and analysis. These models are complemented by a suite of utilities that enhance user experience and functionality. For instance, the platform provides interactive visualization tools for protein embeddings, capabilities for training custom classifiers, and features for conducting empirically scored docking. To facilitate ease of use, these diverse tools and utilities have been encapsulated within a command-line interface, organized into ten commands. This modular approach allows users to effortlessly switch between different approaches, thereby enabling seamless experimentation.

One of the most significant challenges in leveraging cutting-edge models is the initial setup, which often involves configuring a suitable computational environment, installing requisite software packages, and managing intricate dependencies. TRILL streamlines this process, offering an installation procedure that is both straightforward and efficient. By establishing a cache in the end-users home directory, TRILL also intelligently manages downloads to optimize storage usage, where models are downloaded once and only once when first used. Moreover, TRILL addresses the hardware limitations that frequently hinder the effective utilization of LLMs, which can range from millions to billions of parameters. The framework employs advanced techniques such as model parameter sharding to optimize computational resources. Another strength of the platform lies in its flexibility. TRILL’s scalability is a hallmark feature, designed to function across a spectrum of computational environments—from Google Colab and personal laptops without GPU support to high-performance supercomputing clusters. The framework is highly accessible, requiring only protein sequences or structures as user input. TRILL also exposes a range of hyperparameters that can be adjusted to meet specific research needs. Through abstraction, TRILL is able to reduce lines of code needed to be written by end-users by a factor of 10-100.

TRILL’s adaptability is underpinned by robust deep-learning frameworks such as PyTorch Lightning [35] and Hugging Face’s Accelerate [36]. These frameworks facilitate scaling of both training and inference across diverse hardware architectures. We have meticulously transformed many of the neural networks within TRILL into Lightning modules, even going so far as to modify the model’s source code on GitHub. This conversion not only democratizes the models through features like parameter offloading, memory efficient optimizers, and mixed-precision training, but it also facilitates high-performance, distributed analyses on a supercomputer, with a SLURM script example included in the documentation. Additionally, we have engineered all requisite models to operate independently of GPU support. Beyond its core functionalities, TRILL offers auxiliary features like advanced visualization capabilities for protein embeddings. Utilizing dimensionality reduction techniques followed by Bokeh, we generate interactive, queryable scatter-plots that are accessible through a text-input interface powered by Javascript. The platform also leverages key packages such as PyTorch [37], Deepspeed [38], and Transformers [39], along with specialized packages like fair-esm and RFDiffusion [23] for interfacing with relevant deep-learning models.

In the following sections of this manuscript, we will describe two use-cases that demonstrate how TRILL can be employed to compose bespoke workflows. These examples are by no means exhaustive but are intended to inspire further exploration. We believe that the scientific community is just scratching the surface of what is possible when ground-breaking models are chained together into complex pipelines.

## Example TRILL workflows

### Workflow 1: Remote Homology Detection

Remote homology detection with proteins is a complex yet indispensable task in bioinformatics. The importance of identifying distant homologues lies in enabling accurate functional annotation, structural prediction, and evolutionary analysis of proteins, which are fundamental to advancements in drug discovery and understanding diseases [40]. The challenge primarily stems from the need to accurately extract evolutionary information into profiles for effective detection. Traditional sequence-based methods are often inadequate for identifying remote homologues, necessitating more advanced computational techniques like profile-based protein representation. This task holds significant importance for both basic research, such as molecular evolution and protein attribute prediction, and practical applications like 3D protein structure modeling for drug development.

PLMs provide a unique paradigm for remote homology detection that does not rely on sequence alignment. Sequences are first embedded within the high-dimensional latent space of an PLM and then homology detection can be performed by training a classifier to detect sequences from a protein family of interest against a background of out of class sequences. When it comes to viral proteins, the challenges of finding distant relatives are further amplified due to various factors such as rapid evolution, the absence of a single common ancestor, horizontal gene transfer, and overlapping genes. These factors contribute to substantial sequence divergence over short timescales, making viral sequences significantly different from their ancestral forms and unrecognizable to traditional methods like BLAST. A clear example of the sensitivity of these methods can be found in Kirchberger 2022, where they were able to vastly expand the known genomes of *Microviridae* by several thousand through the use of iterative HMM searches for the microviral major capsid protein, or VP1, which is a highly conserved hallmark gene of the viral family [41]. Despite the success of this iterative approach, it required months of searches and thousands of CPU hours. Language model based homology detection can have powerful computational advantages over conventional detection strategies due to the alignment-free nature that facilitates rapid classification. While protein language models such as ESM2 have not been extensively trained on viral sequences, they can still learn a mapping from sequence space to high-dimensional space that might generalize across the tree of life. We apply TRILL for detecting *Microviridae* with a relatively simple workflow, where we first use ESM2 to embed known viral sequences. We then train a classifier and use it to find other VP1s in our simulated pool of viral sequences.

To simulate the viral pool, we downloaded all viral proteins from RefSeq, and then filtered out existing VP1s from the database using an HMM search. To reduce the redundancy, we then performed clustering with MMSeqs2 [40], to bring the number to 130K proteins. After embedding the sequences with ESM2_3B, we trained two types of classifiers for microvirus detection: an XGBoost binary classifier and an Isolation Forest anomaly detector. Our reasoning for including two different types of algorithms lies in the fact that “negative” examples for protein classification are often ill-defined and not experimentally validated. In the case where there is no clear “negative” classification, an anomaly detector just requires one label. We trained the two types of classifiers on a held-out 25% of the VP1 sequences, with the XGBoost also requiring a comparable amount of negative examples for training. We randomly sampled the negative cases for training the binary classifier from the 130K unrelated viral proteins without replacement. After training, the XGBoost and iForests achieved best F1-scores of 0.999 and 0.950 respectively (Figure 3). Remarkably, both classifiers had no false positives, highlighting the utility of these high-dimensional representations for performing sensitive searches.

### Workflow 2: Family Based Protein Generation

Proteins that share similar functions often share ancestry, as they are likely to have evolved from a common ancestral protein. This shared ancestry is usually reflected in sequence similarities, which can be detected through various bioinformatics methods like sequence alignment and profile-based homology detection. However, this is not always the case due to the phenomenon of convergent evolution, where proteins from different evolutionary lineages evolve similar functions independently. In such cases, the proteins may have similar functional domains but differ significantly in their overall sequences, making traditional methods of homology detection less effective. Grouping proteins that share similar functions can be a powerful strategy for fine-tuning generative models. By focusing on these “families”, the model can learn the specific sequence patterns and structural motifs that are crucial for a particular function. This targeted learning can improve the model’s ability to generate novel proteins with desired functionalities, thereby enhancing its applicability in synthetic biology and drug development. Leveraging the knowledge of functionally similar protein families can thus provide a more nuanced and effective approach to protein design.

We apply TRILL to generate novel proteins from a desired grouping. While there are many possible ways to approach this task, our workflow first involves training a functional classifier on ESM2 embeddings from validated positive and negative examples of our protein family. We then proceeded to finetune a generative PLM, ProtGPT2, on known examples of our desired family and compare the classification rates for generated proteins from both the finetuned and base model. For this general workflow, we decided to test it on two different groupings of proteins, cell penetrating peptides and anti-CRISPR proteins.

**Listing 1:**
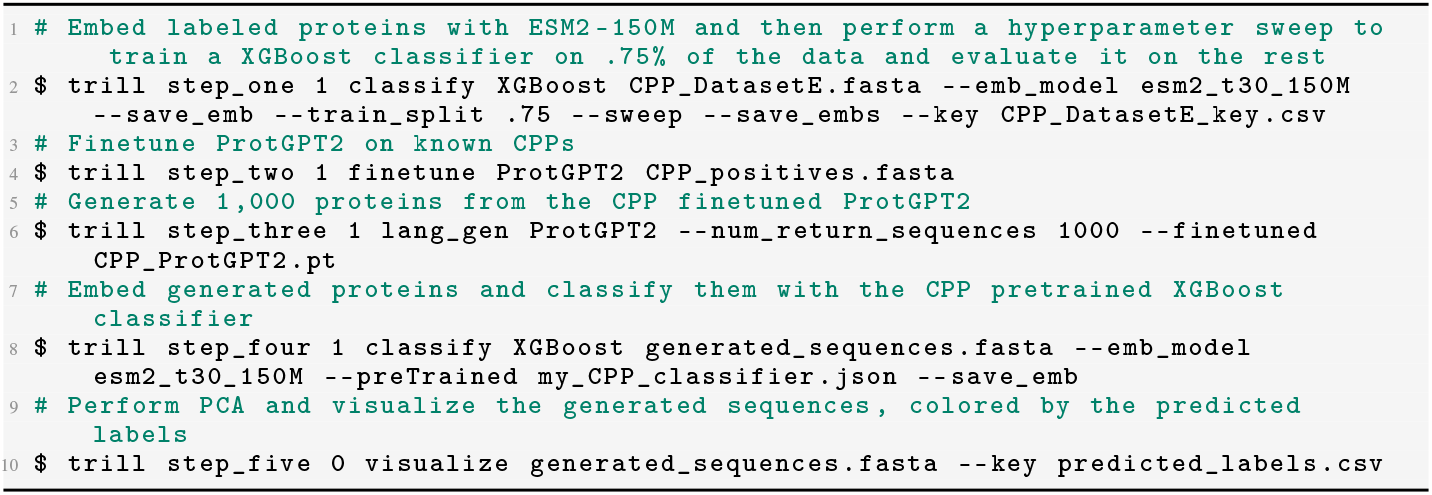
Example TRILL commands for CPP workflow

Cell penetrability is an example of a protein function that BLAST/HMMs routinely fail in identifying due to convergent properties without sharing common ancestry. Utilizing Dataset E from Yadahalli 2020 [8], we first trained an XGBoost classifier on the protein embeddings from ESM2-150M and then achieved an F1 of 0.876 on a held-out 25% of the CPPs (Figure 5). We then finetuned ProtGPT2 on the 955 CPPs for 10 epochs with a learning rate of 1e *−* 5. Next, we generated 1,000 proteins using the base and CPP finetuned ProtGPT2 for each of the top 9 most frequent starting residues used as seed sequences (‘R’: 175, ‘K’: 137, ‘G’: 119, ‘A’: 66, ‘L’: 61, ‘M’: 50, ‘C’: 48, ‘Y’: 44, ‘S’: 38).

We then evaluate the putative de-novo proteins with our pretrained XGBoost classifier. While the base ProtGPT2 model has a non-trivial success rate of generating predicted CPPs, there is a noticeable increase in success with the finetuned model. Between the proteins generated from the base and finetuned models, we predicted an average success rate of 22% and 57% respectively over 9,000 synthetic proteins from both (Figure 2). We surmise that the base ProtGPT2 model might have an artificially high “success rate” due to short peptide length as well as starting with certain amino acids such as arginine and lysine, which both have positively charged side-chains and are the two most frequent starting residues. For further comparison of the generative abilities of these models, we generated two groups of random proteins that have lengths that are drawn from a Poisson distribution (*λ* = 55). The first group is 9000 purely random proteins from a uniform distribution across the 20 canonical amino acids and the second group is 9000 proteins that have a bias to start with a certain amino acid according to the initial residue distribution in the training CPP set, while the rest of the generated sequence is purely random.

**Figure 1:**
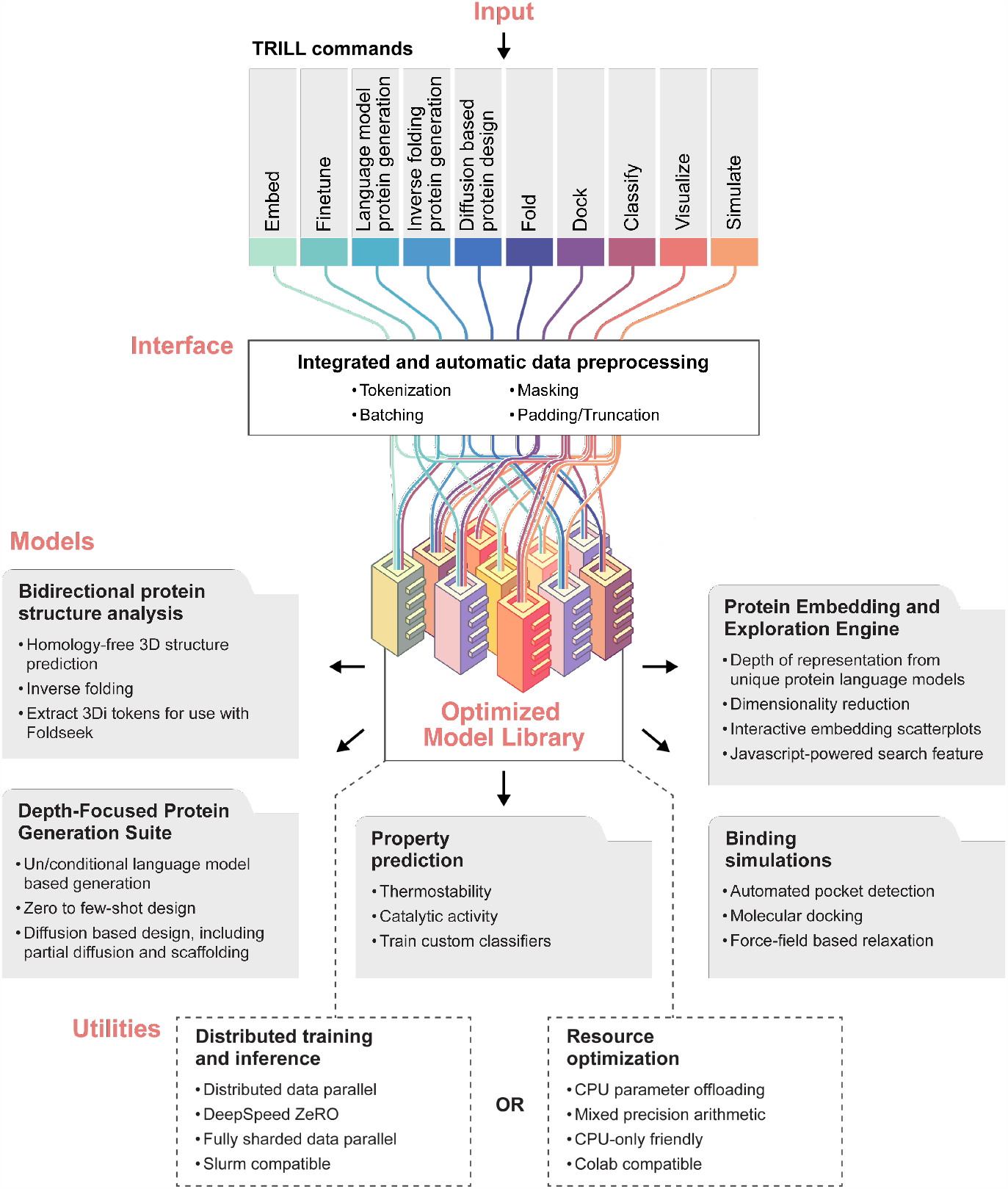
Abstracted TRILL architecture.

**Figure 2:**
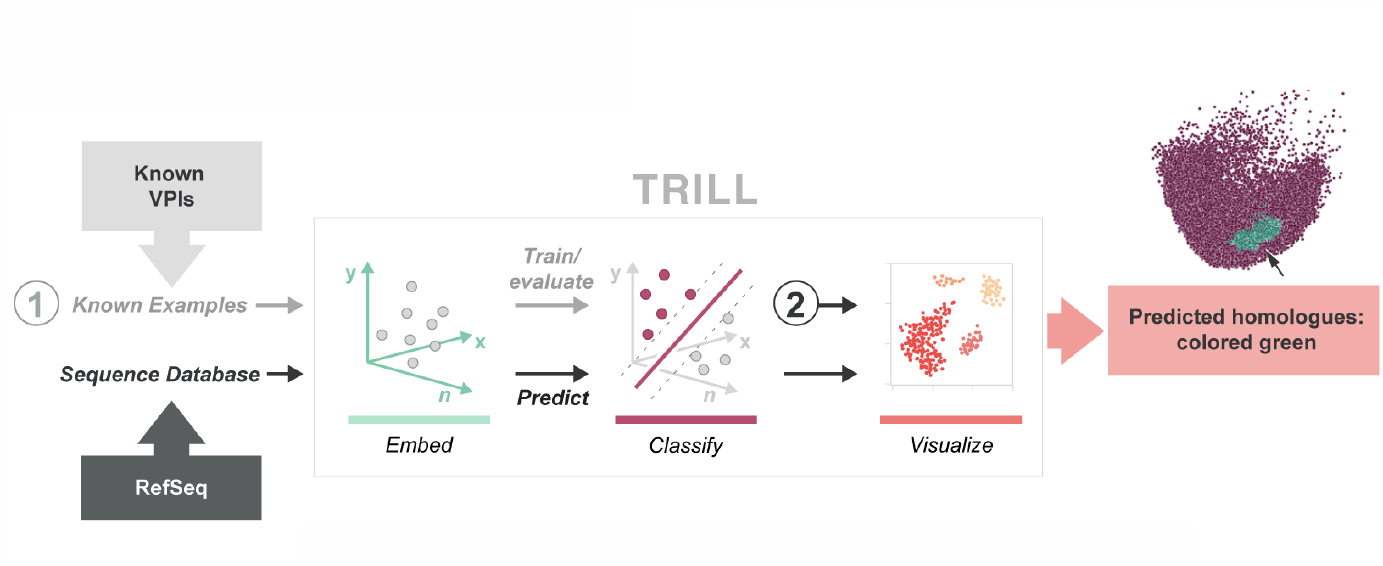
Workflow for finding remote homologues using TRILL

**Figure 3:**
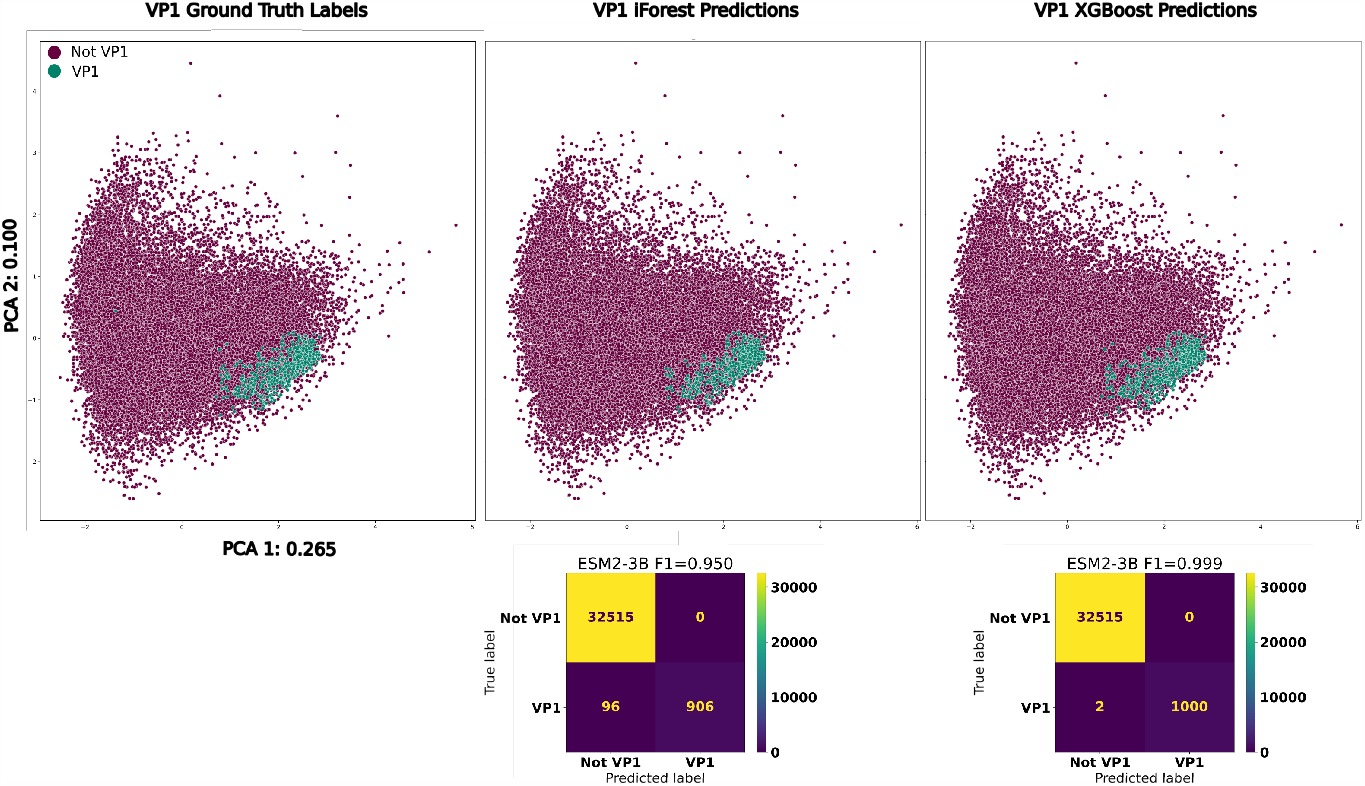
**Top**: PCA of *Microviridae* VP1s, colored by ground truth, iForest and XGBoost predictions respectively from left to right. **Bottom**: Confusion matrices for the iForest and XGBoost VP1 classifiers.

**Figure 4:**
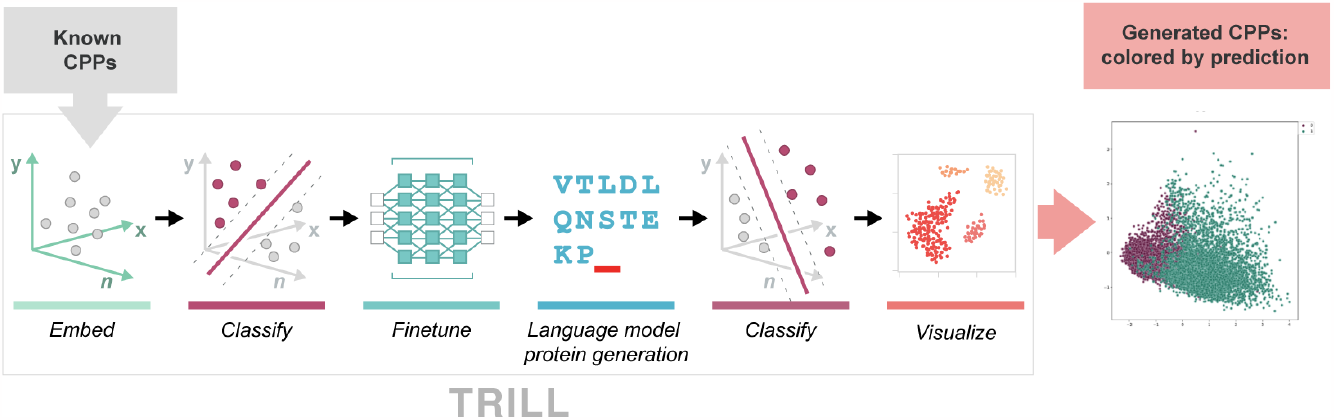
Workflow for generating putative cell-penetrating peptides using TRILL

Anti-CRISPR (ACr) proteins are of great interest to both basic and applied biology due to their ability to inhibit different classes of CRISPR-Cas systems. However, they are also difficult to find in sequencing data due to convergent evolution. While several tools [42, 43] exist for predicting whether or not a protein is an anti-CRISPR protein, we use TRILL to demonstrate how simple it is to train competitive classifiers using nothing but protein embeddings. Leveraging the 150M parameter ESM-2 model, we embedded an equal amount of validated anti-CRISPR proteins and non-anti-CRISPR proteins from PaCRISPR [42]. We then trained an XGBoost classifier on 75% of the embeddings and evaluated the F1 on the held-out 25%, resulting in a F1 score of 0.886. To augment the amount of putative anti-CRISPR proteins, we finetuned ProtGPT2 on 3 different Anti-CRISPR families (I-D, I-F, II-C) for 10 epochs with a learning rate of 1e − 5 and then generated 1,000 sequences from each model, including the base. Quite different from the CPP results, the base ProtGPT2 model only had a predicted 2.1% success rate while the finetuned models achieved 23.5%, 28.7% and 15.7% for I-D, I-F and II-C respectively (Figure 6).

**Figure 5:**
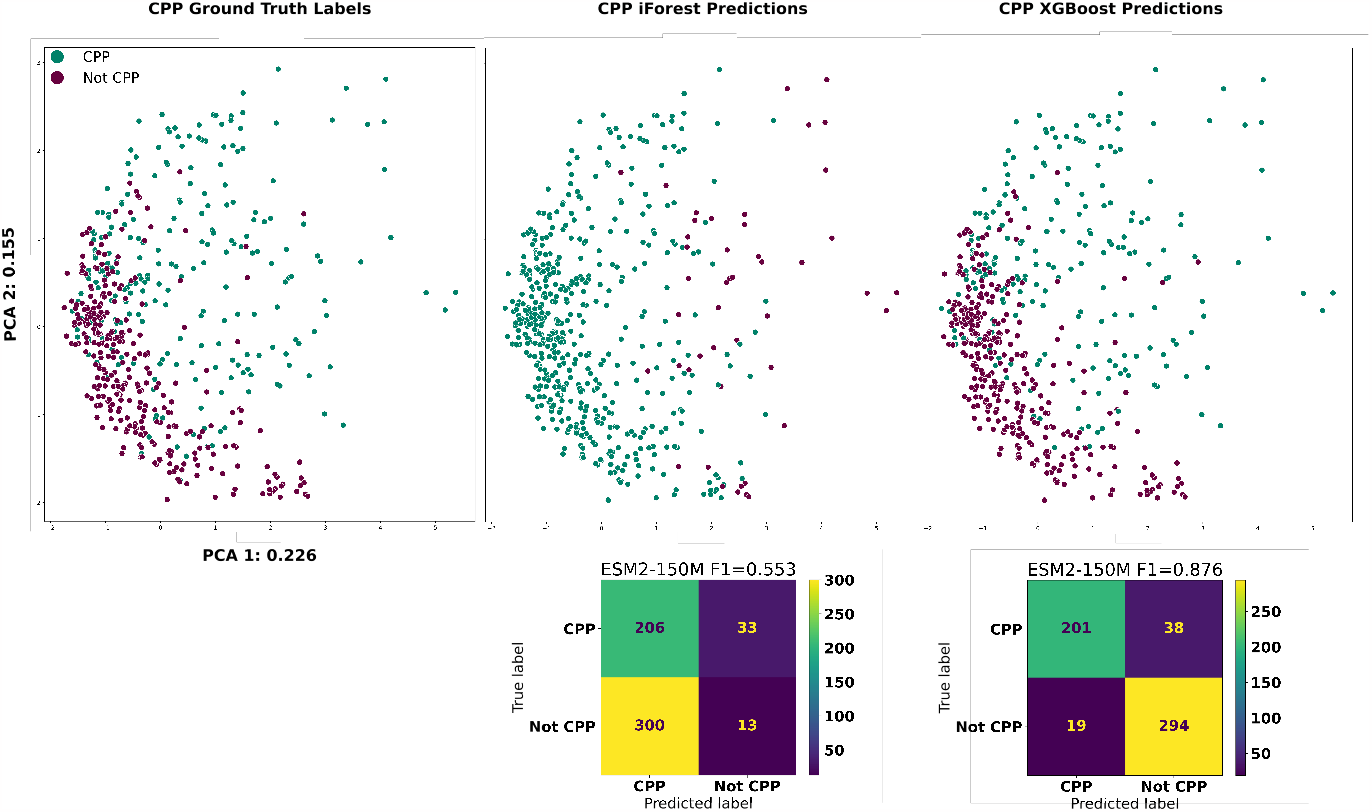
**Top**: PCA of cell penetrating peptides, colored by ground truth, iForest and XGBoost predictions respectively from left to right. **Bottom**: Confusion matrices for the iForest and XGBoost CPP classifiers.

**Figure 6:**
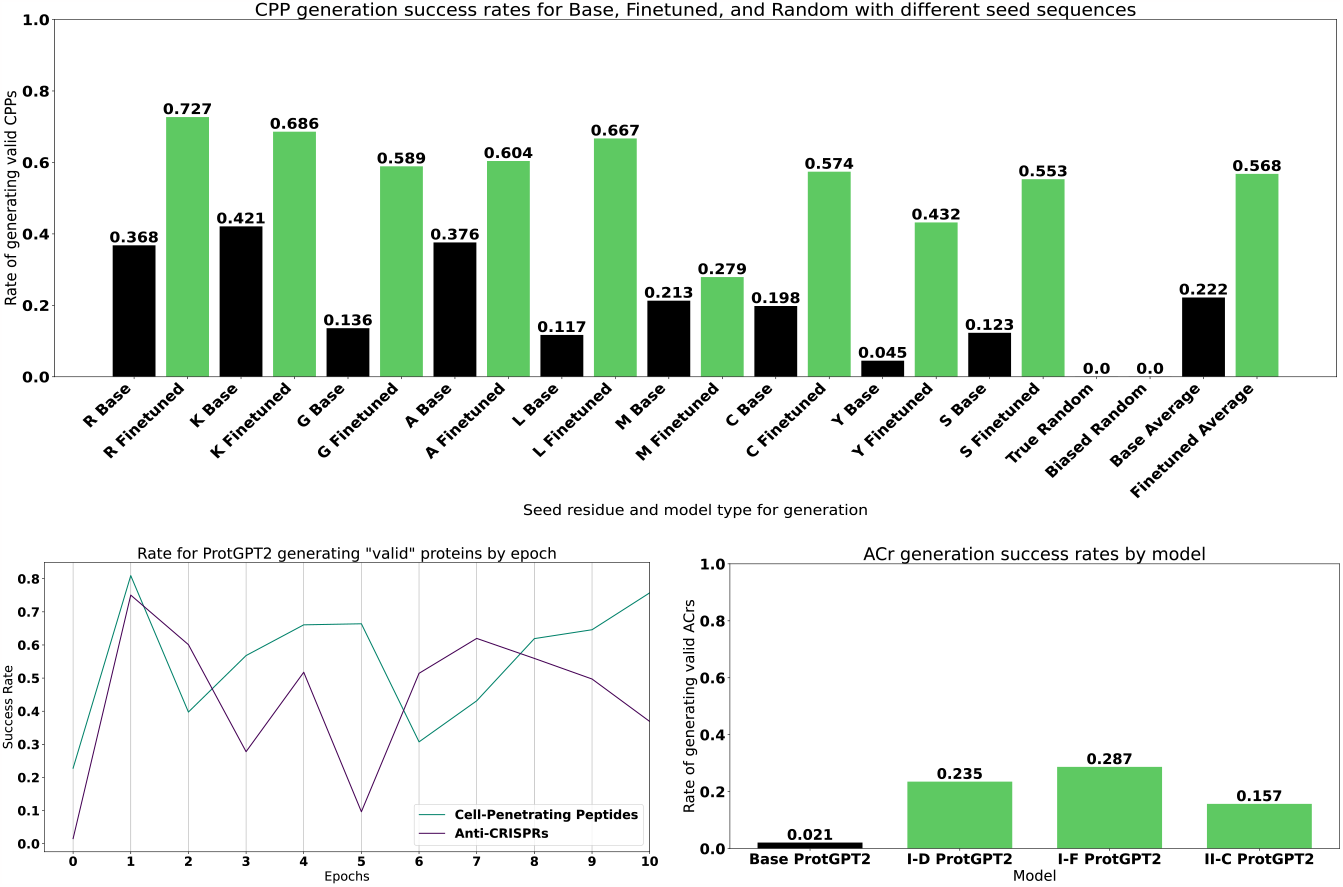
**Top**: Success rates for proteins generated either randomly or from ProtGPT2. In black are the proteins generated from the base model, while the green bars are from the CPP finetuned ProtGPT2. **Bottom Left**: Success rates over the course of finetuning ProtGPT2 for multiple epochs. **Bottom Right**: Success rates for proteins generated from either the base or ACr family specific finetuned ProtGPT2 models.

For both functional classes (CPPs and ACrs), we evaluated how the rate of generating predicted valid proteins varies as a function of finetuning epochs. After each epoch, we generated 5000 proteins and subsequently tested them with our pretrained classifiers. Interestingly, both generative models hit their peak success rate after only 1 epoch, with 95% and 90% respectively for ACrs and CPPs. After 2 epochs, both generative models appear to share similar dynamics, with drops in performance until eventually coming close to the first epoch’s performance. Over the course of 10 epochs, the rate of generating valid proteins varies by around 40% for both models, which highlight the importance of generating proteins from multiple versions of finetuned models. Furthermore, more work is needed to evaluate how the “successful” proteins from 1 epoch of training versus 10 epochs differ. For example, the generative models might end up memorizing certain sub-sequences if finetuned for too many epochs, where the generated proteins are quite similar to the initial training set, potentially losing novel, dissimilar proteins that share function.

### Performance of Classifiers

We systematically evaluated the performance of both XGBoost and iForest classifiers trained on different representations extracted from various protein language models, in order to compare the amount of classification-relevant information that can be learned from the representations. TRILL currently has 3 different types of PLM’s: BERT-style encoders (ESM2), T5-style encoder-decoders (ProtT5, Ankh) and a “bilingual” T5-style model that has been trained on both sequence and structure (ProstT5). Further research is needed to determine which representation is most useful for what application, mainly due to the fact that most usage of these models tends to be for benchmarking purposes so far. Ranging in model size from 8 million to 3 billion parameters, there appears to be diminishing returns as the protein language models grow (Table 2). Interestingly, although the top performing representations are mostly from models with over 1 billion parameters, the smallest model ESM2-8M achieved the highest F1 score in the ACr XGBoost binary classification. Depending on the end-goal, researchers could opt to save on computation and resources by leveraging smaller protein language models for classification. Even though the XGBoost classifiers consistently outperformed the iForests, further comparison of these algorithms is needed, since there seems to be a discrepency in the iForest being able to actually learn protein properties. While it is clear that the iForest can detect remote, ancestrally derived homologies in the case of *Microviridae* VP1’s, the model struggles to identify purely functional classifications, specifically in the case of CPP’s, where the average F1 was 0.582 (Table 2). This disparity in performance might lie in the fact that the CPPs are not ancestrally related, while the ACr’s can be and the VP1’s are, leading to a trade-off between the classifiers when it comes to sequence similarity. There may be some utility in the sequence-conservative predictions of the iForest, but more studies are needed to probe the differences.

**Table 1:** Breakdown of TRILL’s commands.

| COMMAND | FUNCTION |
| --- | --- |
| **Embed** | These models generate numerical representations, or "embeddings," of protein sequences. This transformation allows for quantitative analysis and comparison of proteins, which is crucial for understanding their functions and interactions.<br>*Available models:* ESM2 [11], Ankh [15], ProtT5-XL [16] and ProstT5 [17] . |
| **Visualize** | Performs dimensionality reduction and generates an interactive 2D visualization of the embeddings. Exported as searchable HTML Bokeh scatterplots, this feature powers exploratory data analysis with label-based queries via JavaScript.<br>*Available methods:* PCA [18], t-SNE [19] and UMAP [20] |
| **Finetune** | Finetune protein language models on a specific task with several different training strategies including Distributed Data Parallel and ZeRO-Offload.<br>*Available models:* ESM2, ProtGPT2 [21] and ZymCTRL [22]. |
| **Language Model Protein Generation** | Utilizing pretrained large protein language models, one can either directly generate proteins in a zero-shot fashion or first finetune the models on proteins of interest and then proceed with generation.<br>*Available models:* ESM2, ProtGPT2 and ZymCTRL. |
| **Diffusion Based Protein Generation** | Denosing diffusion probabilistic models have learned to create proteins from atomic noise.<br>*Available models:* RFDiffusion [23]. |
| **Inverse Fold** | Inverse folding is the process of designing a protein sequence that will fold into a desired 3D structure.<br>*Available models:* ESM-IF1 [24], ProteinMPNN [13] and ProstT5. |
| **Fold** | Predict the three-dimensional structure or 3Di tokens (for use with Foldseek [25]) from protein sequence alone.<br>*Available models:* ESMFold [11] and ProstT5. |
| **Dock** | Predicts preferred binding orientations of ligands to a protein.<br>*Available models:* DiffDock [26], Autodock Vina [27], Smina [28] and LightDock [29]. |
| **Simulate** | Performs molecular dynamics simulations to relax protein structures.<br>*Available methods:* OpenMM [30] |
| **Classify** | Enables high-throughput protein property predictions. Users can choose between using pre-trained models or train custom XGBoost [31] and Isolation Forest [32] classifiers on protein embeddings.<br>*Available pre-trained classifiers:* TemStaPro [33] and EpHod [34] |

**Table 2:** Classifier performances.

| PLM | Parameters | Embedding<br>Dimension | CPP<br>XG F1 | CPP<br>iF F1 | ACr<br>XG F1 | ACr<br>iF F1 | VP1<br>XG F1 | VP1<br>iF F1 | Multi-Class<br>Micro F1 |
| --- | --- | --- | --- | --- | --- | --- | --- | --- | --- |
| ESM2-8M | 8M | 320 | 0.854 | 0.546 | 0.916 | 0.733 | 0.978 | 0.833 | 0.830 |
| ESM2-35M | 35M | 480 | 0.853 | 0.563 | 0.895 | 0.714 | 0.980 | 0.862 | 0.835 |
| ESM2-150M | 150M | 640 | 0.876 | 0.553 | 0.886 | 0.745 | 0.984 | 0.906 | 0.842 |
| Ankh | 450M | 768 | 0.839 | 0.613 | 0.888 | 0.751 | 0.993 | 0.956 | 0.826 |
| ESM2-650M | 650M | 1,280 | 0.870 | 0.582 | 0.904 | 0.712 | 0.996 | 0.970 | 0.842 |
| Ankh-Large | 1.15B | 1,536 | 0.833 | 0.601 | 0.882 | 0.773 | 0.986 | 0.443 | 0.813 |
| ESM2-3B | 3B | 2,560 | 0.860 | 0.599 | 0.905 | 0.763 | 0.999 | 0.950 | 0.850 |
| ProtT5-XL | 3B | 1,024 | 0.880 | 0.578 | 0.870 | 0.764 | 0.999 | 0.972 | 0.835 |
| ProstT5 | 3B | 1,024 | 0.886 | 0.609 | 0.889 | 0.802 | 0.998 | 0.689 | 0.826 |
| ESM2-15B | 15B | 5,120 | 0.886 | 0.611 | 0.903 | 0.763 | 1 | 0.963 | 0.844 |

Furthermore, we trained and evaluated XGBoost multi-class classifiers on the three classes of interest (VP1, CPP and ACr), as well as the three “negative” classes. Even though the “negative” classes are not well defined, the classifiers are still able to distinguish between non-cell-penetrating peptides, non-VP1 random viral genes from RefSeq and the non-anti-CRISPR proteins. Focusing on the highest performing classifier for both micro and macro-averaged F1 scores that was trained on ESM2-3B embeddings, we were able to achieve 0.848 and 0.850 respectively (Figure 7). However, all the representations, ranging from 320 to 2,560 dimensions, perform comparably in the multi-class setting, with only 0.018 and 0.02 differences in micro and macro-averaged F1 specifically between the largest, most successful model ESM2-3B and the smallest, ESM2-8M (Table 2). From the ROC curves, it is apparent that the classifiers are robust to false positives, with AUC between 0.81 and 1.00 (Figure 7). Lastly, the multi-class confusion matrix highlights where the classifier had difficulty distinguishing between non-anti-CRISPR proteins and non-VP1 viral genes. This most likely has to do with the fact that these two classes are not mutually exclusive and could definitely have overlap, since the anti-CRISPR negative examples come from phages and mobile genetic elements in bacteria [42].

**Figure 7:**
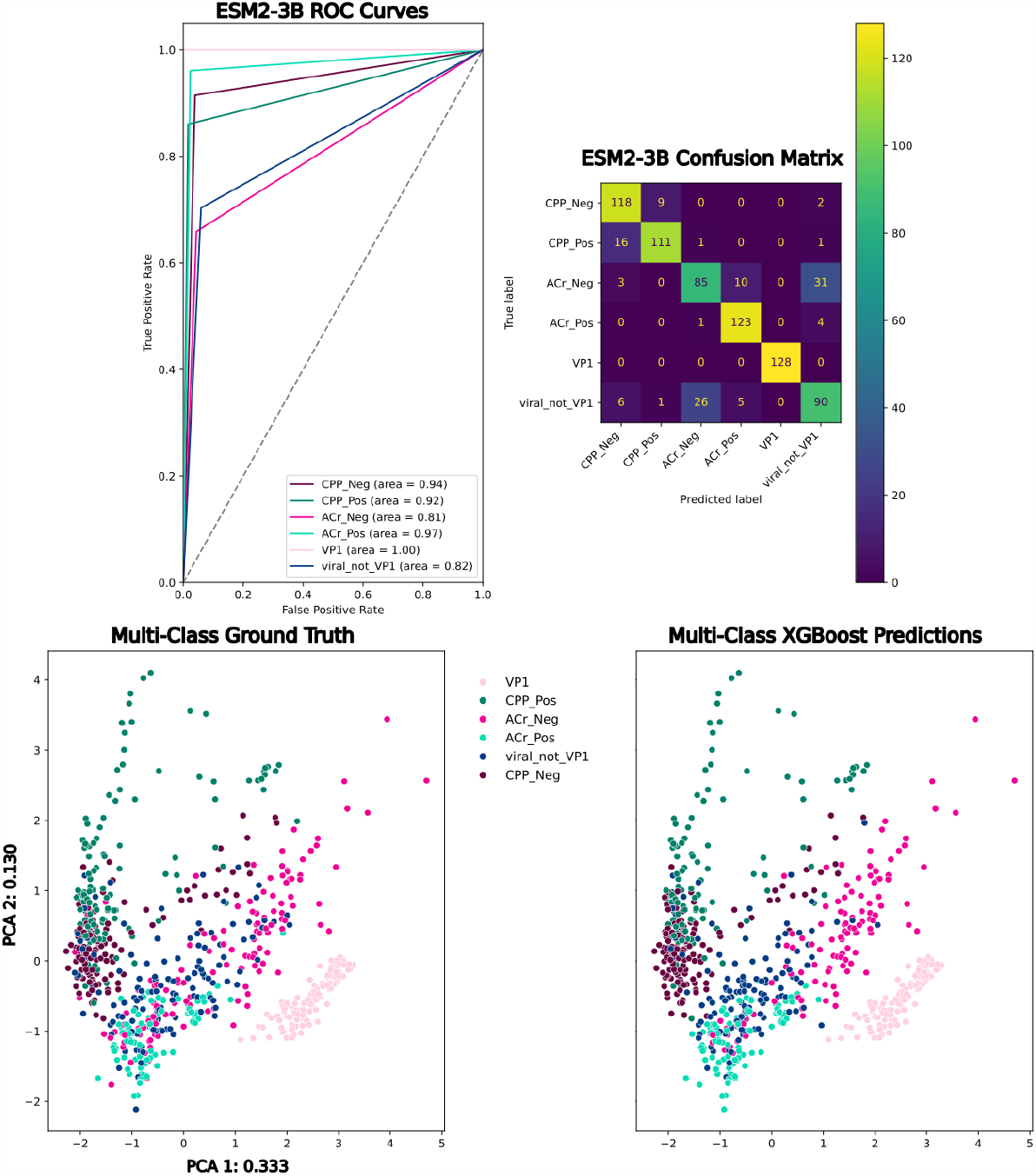
**Top Left**: One-versus-all ROC curves from an XGBoost classifier trained on 6 classes represented by ESM2-3B. **Top Right**: Multi-class confusion matrix for all 6 classes represented by ESM2-3B **Bottom Left**: PCA of ESM2-3B embeddings colored by ground-truth labels. **Bottom Right**: PCA of ESM2-3B embeddings colored by predicted labels.

## Conclusion

The workflows presented in this manuscript are intended as illustrative examples to stimulate further inquiry; they should not be viewed as an exhaustive catalog of TRILL’s potential applications. As TRILL undergoes iterative advancements, the incorporation of cutting-edge deep-learning models serves to diversify the range of scientific inquiries that can be addressed. TRILL has only been made possible through building upon the efforts of open-source software development, highlighting the importance of open-science and collaboration. As the field of deep-learning-based bioengineering matures, the real-world impact won’t be fully realized without the sharing of methods and widespread adoption. With broader use of these tools, an immediate benefit is the empirical validation of these integrated algorithms through rigorous experiments performed independently. Such validation is poised to bolster the scientific community’s confidence in these methods and provide invaluable data for subsequent algorithmic refinements. A promising direction for future development is the implementation of specialized, end-to-end declarative pipelines. These automated pipelines would allow end-users to execute complex analyses with minimal manual intervention, thereby accelerating the pace of discovery. The workflows will be designed to strike a balance between generalizability and customizability, thereby accommodating a broad spectrum of bioengineering challenges while also allowing for task-specific configurations.

In conclusion, TRILL aims to be a facilitative platform, offering a new lens through which to approach proteins. Its potential impact, while promising, will ultimately be determined by its ability to integrate seamlessly into existing research workflows and to adapt to the rapid pace of both biotechnology and artificial intelligence. It is through this iterative process of development, validation, and adaptation that TRILL aspires to become a meaningful tool for advancing science.

## Data Availability

All of the protein sequences, embeddings, finetuned generative models and other artifacts for this work are available for download at https://doi.org/10.22002/mn4w0-cqj07. The plots in the manuscript and analyses can be reproduced by following the included Jupyter notebooks.

## Acknowledgements

Special thanks to Inna Strazhnik for making very informative figures to showcase TRILL. Also more special thanks to Martin Holmes for bravely testing alpha TRILL, as well as to Arjuna Subramanian and Alec Lourenço for being early adopters and avid users of TRILL, they have provided valuable insights and found manu bugs. Thanks also to Lucas Schaus for the idea of comparing classified CPPs from ProtGPT2 to “junk” random proteins. Another thanks to Changhua Yu for helping me dive into PLMs, with very useful Jupyter notebooks using ESM1 that helped me get started in this fast paced field. The authors would also like to thank Shilpa Yadahalli for cleaning and sharing Dataset E through personal communication. ZAM would also like to thank G Anthony Reina for contributing to the TRILL documentation and Blossom Market Hall in San Gabriel for providing a comfortable setting and free Wi-Fi to work on TRILL. This work was supported by the Heritage Medical Research Institute, Gordon and Betty Moore Foundation, NIH R01-GM150125, and the Packard Foundation.

